# DNAscope: High accuracy small variant calling using machine learning

**DOI:** 10.1101/2022.05.20.492556

**Authors:** Donald Freed, Renke Pan, Haodong Chen, Zhipan Li, Jinnan Hu, Rafael Aldana

**Affiliations:** Sentieon Inc., San Jose, CA

## Abstract

We present DNAscope, an accurate and efficient germline small-variant caller. DNAscope combines the robust and well-established preprocessing and assembly mathematics of the GATK’s HaplotypeCaller with a machine-learned genotyping model. Benchmarks of DNAscope and DNAseq (Sentieon’s GATK-matching germline variant calling pipeline) demonstrate that DNAscope achieves superior SNP and insertion/deletion accuracy with reduced computational cost.

---

The Genome Analysis Toolkit (GATK) HaplotypeCaller has become the industry standard variant small variant caller due to its high accuracy. By combining Bayesian statistical models with direct modeling of read haplotypes and well-tuned variant filters^1^,2HaplotypeCaller has achieved top performance in a variety of public and third-party benchmarks^3–5^. However, existing variant callers based on short read technology including HaplotypeCaller have less than perfect matching to high-confidence variant calls, especially across complex genomic regions such as homopolymers and other repetitive loci. Many of these complex regions are clinically relevant and improving variant calling accuracy at these sites is increasingly important as next-generation sequencing data becomes more frequently used in clinical assays^6,7^.

Most variant callers use expertly constructed statistical models for variant genotyping and implement hand-tuned filters with reasonable default settings to remove false-positive variant calls. While these filters work well in most cases, machine learned filters and genotyping models have the ability to improve accuracy by learning more complex relationships between variant features. Recent improvements in both the quality of genomic reference materials and machine learning methodologies have led to the rise of a number of approaches that move away from explicit statistical models to machine learned models of sequencing error modes that improve variant filtration or genotyping^8,9^.

DNAscope uniquely combines the well-established methods from haplotype-based variant callers with machine learning to achieve improved accuracy. As a successor to GATK HaplotypeCaller, DNAscope uses a similar logical architecture, but introduces improvements to active region detection and local assembly for improved sensitivity and robustness, especially across high-complexity regions. When a machine learning model is applied, DNAscope outputs candidate variants with additional informative annotations. Annotated variant candidates are then passed to a machine learning model for variant genotyping, resulting in improvements in both calling and genotyping accuracy (Fig. 1, Methods). For non-mammalian species, DNAscope can be used with a Bayesian genotyping model, allowing users to benefit from DNAscope’s improved active region detection and local assembly when resequencing diverse organisms.

**Fig 1.**
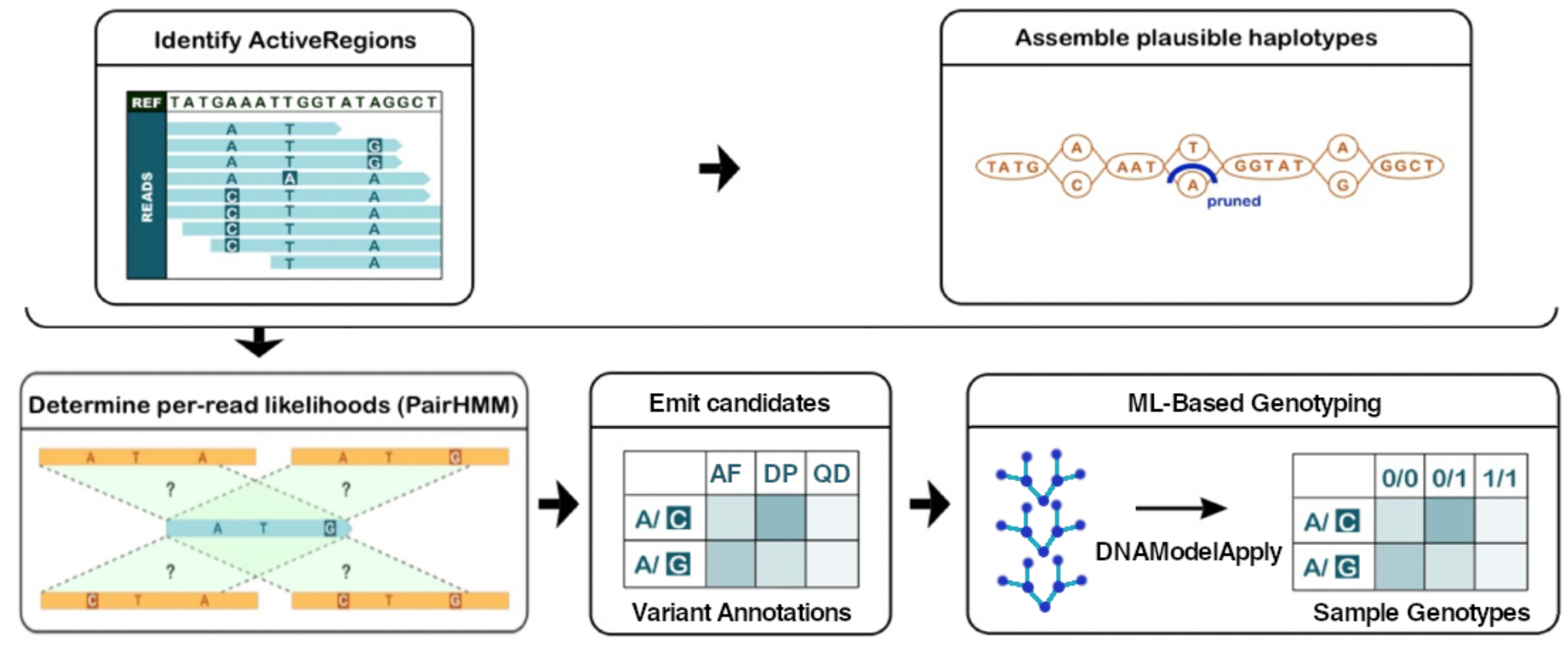
Overview of the DNAscope methodology. DNAscope follows a similar algorithmic flow to GATK HaplotypeCaller. Sites likely to harbor genetic variation are identified as active regions. Sequence reads aligned across active regions undergo local assembly using *de Bruijn* graphs and read-haplotype likelihoods are calculated through PairHMM. Variant candidates are then annotated and emitted. Machine Learning-based genotyping processes variant candidates to determine the correct variant genotype. Modified from a published study^2^.

To assess the variant calling accuracy of DNAscope across a variety of individuals, we called variants with Sentieon’s DNAscope and DNAseq (matching the GATK germline Best Practices^10–12^) pipelines using publicly available data from three GIAB samples: HG002, HG003, and HG004. Samples HG001 and HG005 were not benchmarked as they were used during DNAscope model training. All three benchmarked samples have an approximate coverage of 36x and subsampling was conducted to create five datasets covering depth from 15x – 36x. We benchmarked the performance of these pipelines against the NIST GIAB high-confidence calls v4.2.1 (Fig. 2-5)^5,13^. It is worth noting that the recently updated v4.2.1 truthset includes regions that are challenging for short reads sequencing technologies and our F1-scores are not comparable to benchmarks with older (easier) versions of the GIAB truthset. We also omit variant quality score recalibration (VQSR) as it was reported VQSR significantly decreases the measured F1-score^14^.

**Fig. 2.**
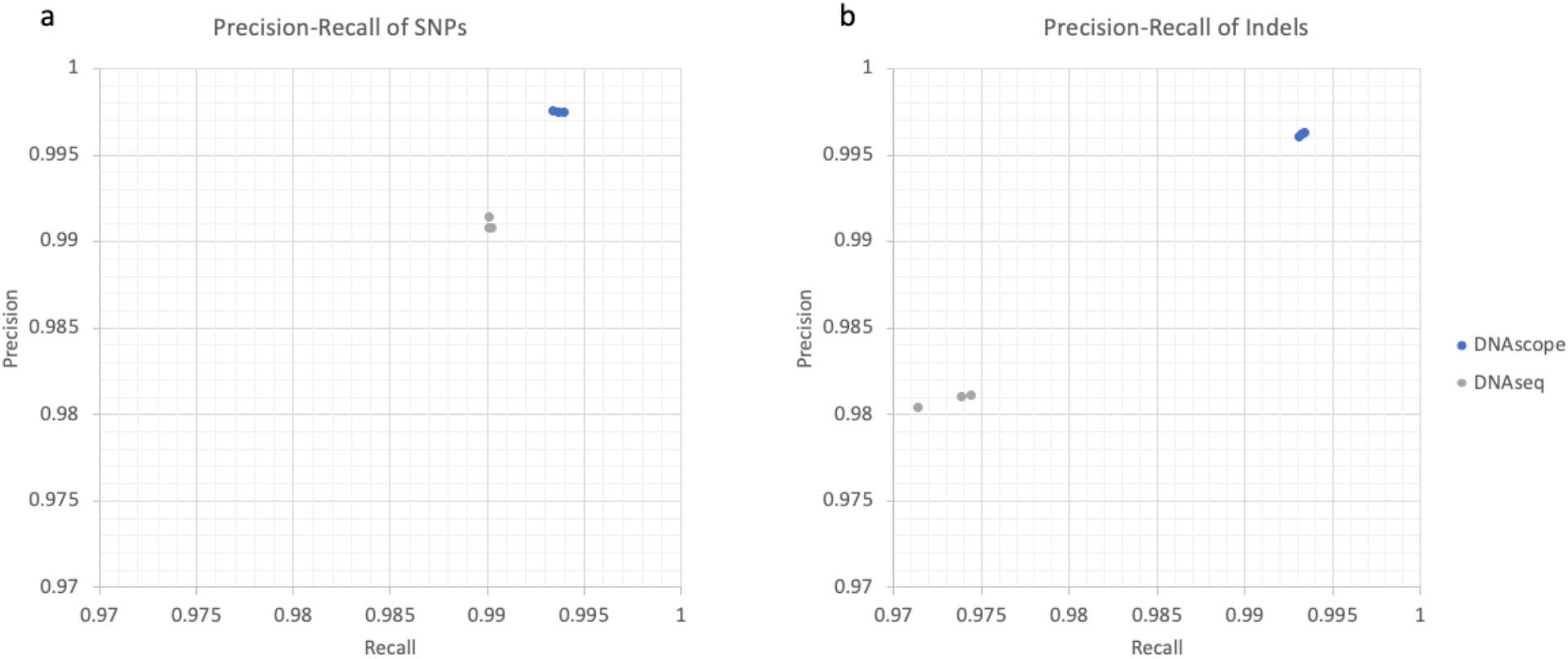
Precision-recall curves of DNAscope and DNAseq. Precision-recall metrics for 30x dataset of sample HG002, HG003, and HG004 for **a**, SNP and **b**, INDEL. DNAscope provides consistently higher accuracy than DNAseq.

For both SNPs and INDELs, DNAscope’s recalls and precisions are significantly higher than DNAseq on all samples at all coverages. For the 30x HG002 sample, DNAscope achieves an F1-score of 99.57% for SNPs and 99.46% for INDELs. Notably, DNAscope reduces total errors by more than two-fold (from 87,607 to 34,634) relative to DNAseq. The high accuracy of DNAscope across multiple samples suggests that DNAscope’s model is not overfit to the HG001 and HG005 samples used during model training. It should also be noted that model training is sequencing platform dependent and an MGI model was developed and benchmarked in a previously published study^15^, demonstrating that DNAscope may be adapted to new sequencing technologies.

To further explore the strengths of the benchmarked variant callers, we stratified the performance of the tools in our benchmark using Global Alliance for Genomics and Health (GA4GH) stratification regions (Fig. 3)^16^. The GA4GH stratification regions define regions of low “mappability”, where accurate alignment of short sequencing reads to the reference genome is difficult due to the presence of multiple, long and highly similar sequences in the reference genome. Additional stratifications highlight other regions such as segmental duplications, self-chain regions (regions of non-trivial alignment of a reference to itself), the major histocompatibility complex (MHC), and “alldifficult” regions, which combine low mappability, long homopolymer, tandem repeats, extreme GC content and other difficult regions. Stratified accuracy analysis finds that DNAscope substantially outperforms DNAseq across stratifications associated with difficulty in read mapping, including the “lowmappabilityall”, “segdups”, and “chainSelf” stratifications (Fig. 3). DNAscope additionally has higher SNP accuracy across the MHC and higher INDEL accuracy across long homopolymers contributing to DNAscope’s better SNP and INDEL performance across difficult regions. Across easier regions, DNAscope and DNAseq have similar SNP accuracy, while DNAscope maintains a higher INDEL accuracy.

**Fig. 3.**
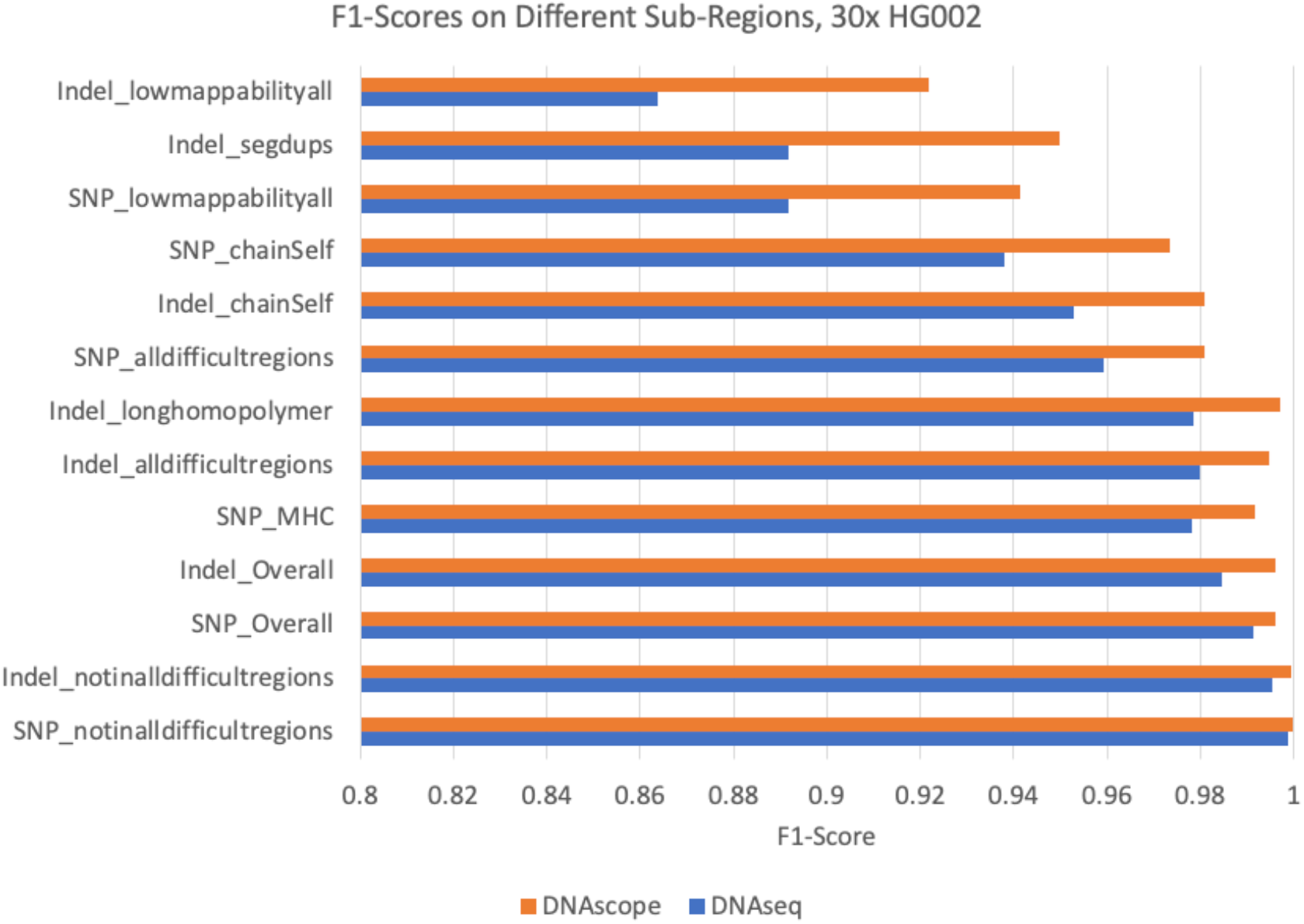
Evaluation across GA4GH stratification regions, HG002 at 30x depth. F1-score when compared against the NIST GiaB high-confidence calls for DNAscope and DNAseq across GA4GH stratification regions for **a**, SNP and **b**, INDEL. The bar chart shows the F1-score for the benchmarked tools (bottom axis). DNAscope has higher accuracy across most stratified regions. Note that stratification regions are not exclusive and one variant can be included in multiple stratified analyses. Stratifications are ordered by the delta between DNAscope and DNAseq, and stratifications with less than 10,000 variants were excluded.

To assess how the benchmarked variant callers’ accuracy changes with sequencing coverage, we set out to study the performance across 5 sequencing depth using subsampled 36x data sequenced from HG002, HG003, and HG004 (Fig 4). Variant calling accuracy typically drops with lower sequencing coverage; however, we noticed DNAscope maintained a higher accuracy at lower coverages compared to DNAseq, indicating variant calling benefits more from DNAscope’s improved architecture and machine learning model filtering at lower coverages. Notably, DNAscope at 20x coverage has consistently higher accuracy than the GATK-matching DNAseq pipeline at 36x coverage.

**Fig. 4.**
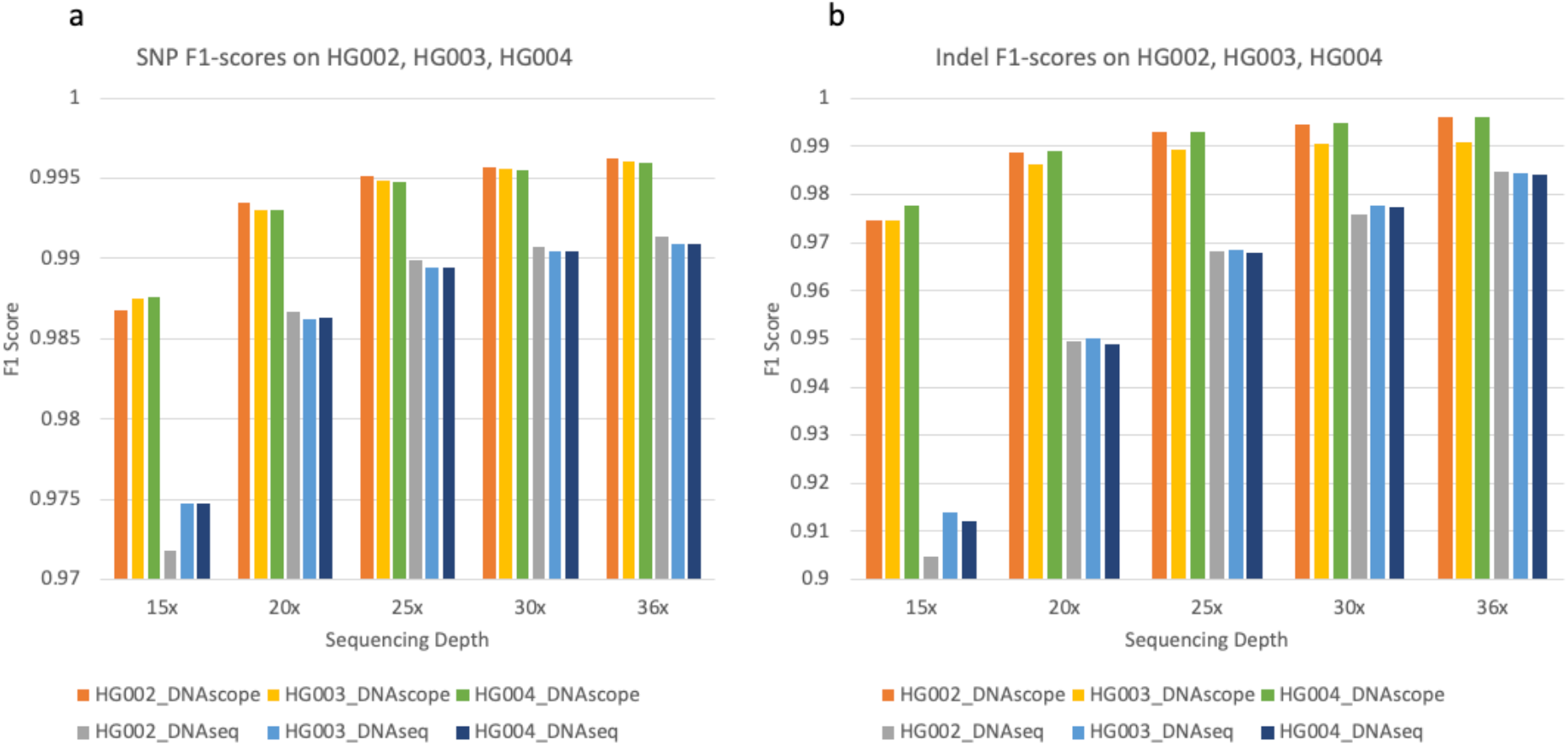
Evaluation on Genome in a Bottle sample HG002, HG003, HG004 with multiple sequencing depths. F1-score measures for DNAscope and DNAseq for SNP (**a**) and INDEL (**b**).

Resequencing of non-human and polyploid samples is common in agricultural genomics. DNAscope’s machine learning model is designed to work with diploid samples and current versions of DNAscope do not support non-diploid samples. Accordingly, we sought to explore the accuracy of DNAscope using DNAscope’s Bayesian genotyping model (Fig 5). We specifically displayed performance of the Bayesian model at 15x coverage, representing the typical coverage of samples with large genomes where high-depth sequencing may not be affordable. As expected, DNAscope’s Bayesian genotyping model decreased overall accuracy compared to the machine-learned genotyping model. While DNAscope’s Bayesian SNP F1-score was similar to DNAseq, DNAscope outperformed DNAseq’s INDEL F1-score.

**Fig. 5.**
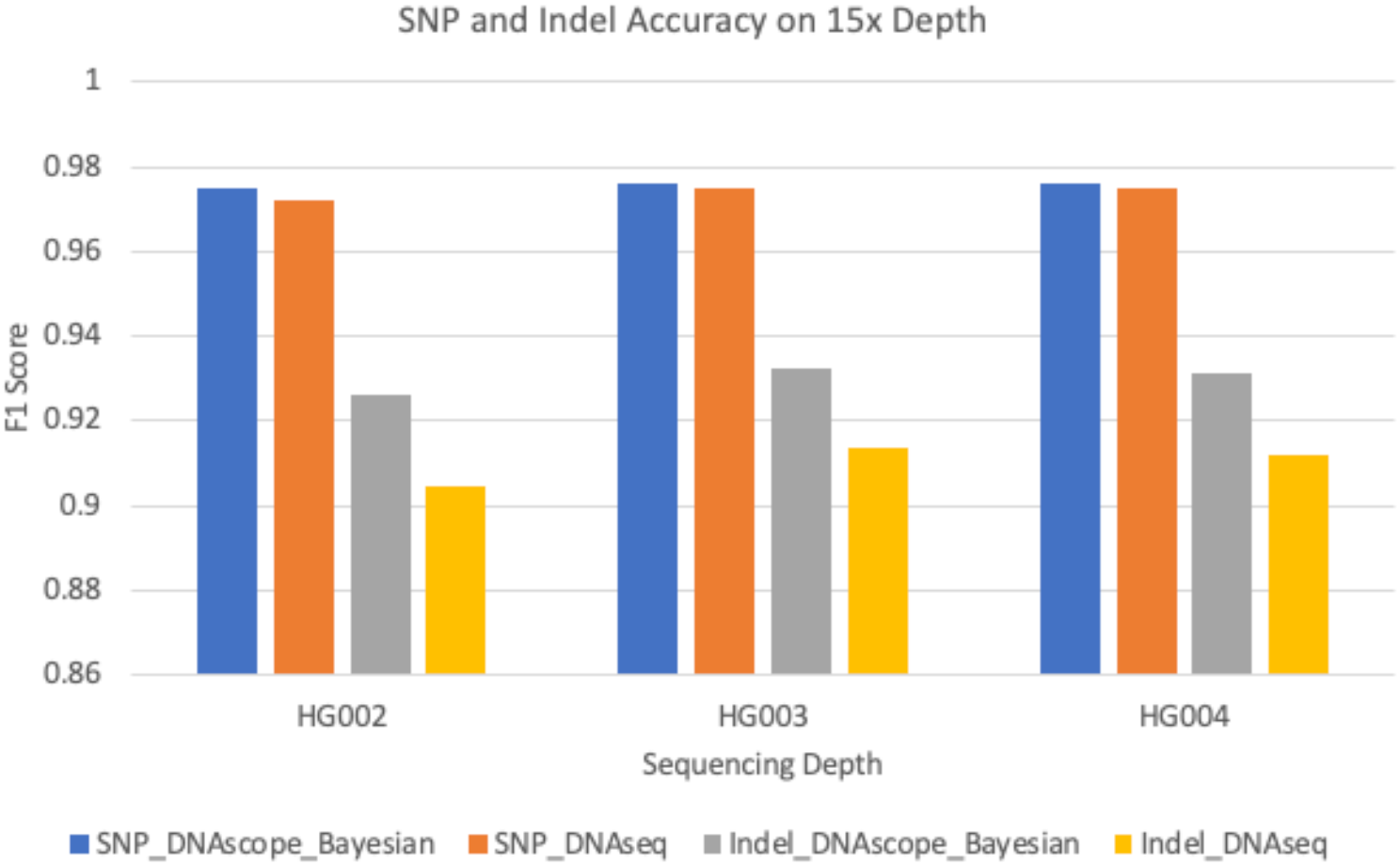
Evaluation on Genome in a Bottle sample HG002, HG003, and HG004 with 15x sequencing depths. F1-score measures for DNAscope’s Bayesian genotyping model and DNAseq for SNP and INDEL.

While not the primary focus of this study, our experimental design also allowed us to benchmark the runtime of DNAscope in a standardized environment. A 30x HG002 dataset was processed on four AWS C6i instances (Intel Xeon CPU at 2.9 GHz). DNAscope’s runtime was plotted by stage. As shown in Fig 6, DNAscope processed a 30x WGS sample in less than 1 hour on machines with more than 96 vCPUs, which is similar to DNAseq’s runtime and approximately five times faster than opensource BWA/GATK pipeline^12^. Good scalability could also be observed, and runtime is almost linearly correlated with the number of available threads.

**Fig. 6.**
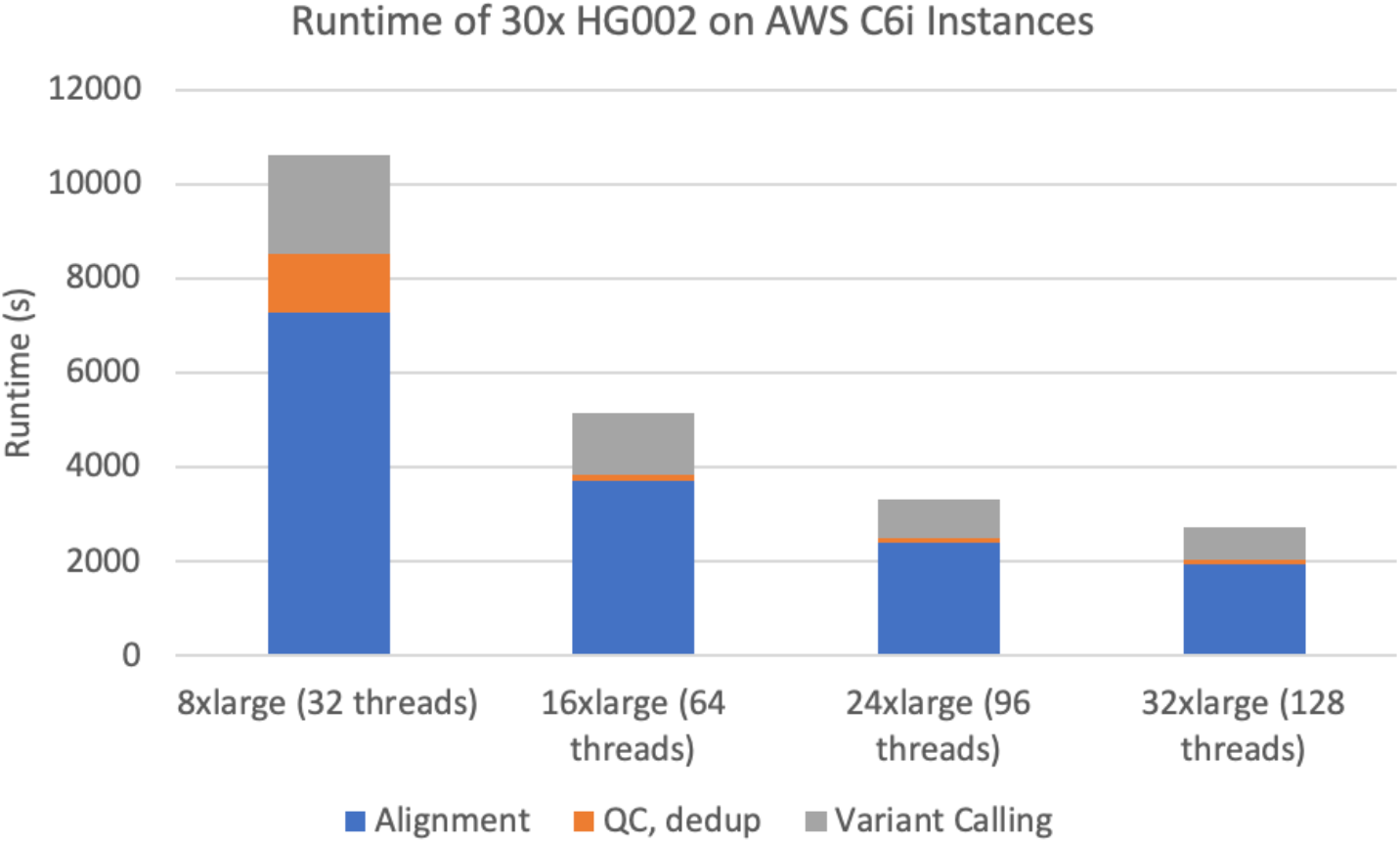
Runtime of DNAscope on multiple AWS C6i Instances. Runtime is plotted per stage for each instance type.

In this work we demonstrate that DNAscope achieves higher accuracy than DNAseq across samples and different levels of coverage. A stratified analysis using GA4GH stratification regions enabled us to confirm the high accuracy of DNAscope across the majority of stratified regions and highlighted DNAscope higher accuracy across indels and stratifications containing genomic regions where variant calling is more difficult. DNAscope combines the well-established mathematical and statistical models used in the GATK’s HaplotypeCaller with machine learning for variant genotyping to achieve superior accuracy while maintaining computational efficiency.

## Author Contributions

D.F., R.P., H.C., and R.A. designed the experiments. D.F., R.P., Z.L, and J.H performed the analysis. D.F., R.P., H.C., and J.H wrote the manuscript with input from all authors.

## Methods

### Improvements in variant candidate detection

#### Overview

Sentieon Haplotyper is a re-implementation of the mathematics of the GATK’s HaplotypeCaller that uses more efficient computational algorithms to achieve speed improvements while providing matching results^10–12^. Both Haplotyper and DNAscope use the same overall architecture as HaplotypeCaller, including active region detection, local assembly, calculation of read-haplotype likelihoods, while improving robustness through removing down-sampling, an improved implementation, better resource management and improved computational algorithms. DNAscope includes additional minor implementation corrections and variant annotations as described below.

#### Variant annotations

DNAscope outputs additional variant annotations that are not used by the GATK. These and the standard annotations are used by the machine learning model during variant genotyping.

“Entropy” is defined as the Shannon entropy of all haplotypes discovered during local assembly. High entropy is an indicator of the presence of mismapped reads or other read alignment issues.

“UDP” is the unfiltered read depth across all input samples. This metric is similar to “DP”, however “DP” removes reads with low mapping quality or a high likelihood of being mismapped.

“NBQ” is the mean base quality of the variant base and 5bp on either side of the variant candidate. The “NBQ” annotation is inspired by the “NBQ” annotation in the Atlas2 Suite variant analysis pipeline^17^.

“ReadPosEndDist” is the mean distance from either end of the read. The “ReadPosEndDist” annotation is inspired by the “ReadPosEndDist” annotation in bcbio-nextgen^18^.

### Model-based genotyping

DNAscope’s genotype model is designed to help distinguish systematic noise from true germline variants. Through machine learning, we aim to capture the error patterns from library prep, sequencing, and alignment as well as to extract informative signals from regions that are inherently difficult to interpret. To achieve improved efficiency while maintaining competitive performance with deep learning approaches, Gradient Boosting Machines (GBMs) are used to learn patterns from the structured data^19^. GBMs build trees in succession to train sequential ensembles of weak, base learners, reducing residuals in a stepwise fashion. The incremental improvement from each base learner is controlled through the learning rate, allowing early stopping if any overfitting is observed during the validation process.

A GMB is trained using data from the HG001 and HG005 GIAB samples from various data sources, while excluding chromosome 20. During model training, 90% of variants are used in the training process while the remaining variants are used as a validation set to optimize the model complexity for the bias/variance tradeoff and prevent overfitting. Input to the model is a DNAscope VCF where DNAscope has been run with sensitive settings, allowing for the detection variant candidates with very little read support. The model then utilizes the annotations from the DNAscope VCF output as features to classify 4 genotypes: “0/0”, “0/1”, “1/1”, “1/2”, assuming the sample to be diploid across all loci. The genotype “0/0” is used to indicate that the reported variant is classified as a reference call instead of a true genetic variant. From the verbose list of variant candidates, the DNAscope genotype model reduces the false discovery rate drastically, while updating sample genotypes and maintaining high sensitivity. In addition, the model also adds the annotation “ML_PROB” to the VCF file, which is the probability of false discovery (the actual genotype is 0/0) as calculated by the model.

### Benchmarking Methodology

Public FASTQ data were obtained from their respective sources as described in the Data availability section. All FASTQ were processed using a standardized data processing pipeline including read alignment with Sentieon improved BWA-MEM version 202112.01, duplicate marking using Sentieon version 202112.01, subsetting using SAMtools version 1.3.1 (applied to some samples), variant calling using DNAscope version 202112.01, DNAscope Illumina model 1.0, Haplotyper 202112.01, and variant evaluation using hap.py version 0.3.10 with RTGtools vcfeval version 3.8.2 as the variant comparison engine. NIST truth set v4.2.1 was used in evaluation. Intermediate and output files produced during analysis can be found at gs://dnascope-benchmarking. Hap.py output and other output files were analyzed using interactive and collaborative Jupyter notebooks on Google’s Colaboratory.

## Code availability

DNAscope is a commercial software developed and marketed by Sentieon Inc., San Jose, CA. Sentieon will provide a free short-term evaluation license that can be used to replicate this work and may be obtained by contacting. The software package used in these experiments can be downloaded from https://s3.amazonaws.com/sentieon-release/software/sentieon-genomics-202112.01.tar.gz, while DNAscope’s machine learning model can be downloaded from https://s3.amazonaws.com/sentieon-release/other/SentieonDNAscopeModel1.0.model.

## Data availability

GIAB data are publicly available from the GIAB ftp site, ftp://ftp-trace.ncbi.nlm.nih.gov/giab/ftp/.

## References

1. Garrison, E. & Marth, G. Haplotype-based variant detection from short-read sequencing. ArXiv12073907 Q-Bio (2012).

2. Poplin, R. et al. Scaling accurate genetic variant discovery to tens of thousands of samples. bioRxiv (2018).

3. PrecisionFDA Truth Challenge – precisionFDA. Available at: https://precision.fda.gov/challenges/truth/results. (Accessed: 10th March 2019)

4. Zook, J. M. et al. Extensive sequencing of seven human genomes to characterize benchmark reference materials. Sci. Data 3, 160025 (2016).

5. Zook, J.M., McDaniel, J., Olson, N.D. et al. An open resource for accurately benchmarking small variant and reference calls. Nat Biotechnol. 37, 561–566 (2019).

6. Goldfeder, R. L. et al. Medical implications of technical accuracy in genome sequencing. Genome Med. 8, 24 (2016).

7. Wagner, J., Olson, N.D., Harris, L. et al. Curated variation benchmarks for challenging medically relevant autosomal genes. Nat Biotechnol. (2022).

8. Poplin, R. et al. A universal SNP and small-indel variant caller using deep neural networks. Nat. Biotechnol. 36, 983–987 (2018).

9. Kim, S. et al. Strelka2: fast and accurate calling of germline and somatic variants. Nat. Methods 15, 591 (2018).

10. Jessica A. Weber, Rafael Aldana, Brendan D. Gallagher & Jeremy S. Edwards. Sentieon DNA pipeline for variant detection - Software-only solution, over 20× faster than GATK 3.3 with identical results. PeerJ Preprints (2016).

11. Freed, D. N., Aldana, R., Weber, J. A. & Edwards, J. S. The Sentieon Genomics Tools - A fast and accurate solution to variant calling from next-generation sequence data. bioRxiv (2017).

12. Kendig, K. et al. Sentieon DNASeq variant calling workflow demonstrates strong computational performance and accuracy. Front. Genet. 20 August (2019).

13. Wagner, J. et al. Benchmarking challenging small variants with linked and long reads. Cell Genomics. Volume 2, Issue 5, 11 May 2022.

14. Zhao, S., Agafonov, O., Azab, A. et al. Accuracy and efficiency of germline variant calling pipelines for human genome data. Sci Rep 10, 20222 (2020).

15. Shen, H. et al. Advanced Whole Genome Sequencing Using an Entirely PCR-free Massively Parallel Sequencing Workflow. bioRxiv (2020).

16. Krusche, P. et al. Best practices for benchmarking germline small-variant calls in human genomes. Nat. Biotechnol. 1 (2019).

17. Challis, D. et al. An integrative variant analysis suite for whole exome next-generation sequencing data. BMC Bioinformatics 13, 8 (2012).

18. Validated, scalable, community developed variant calling, RNA-seq and small RNA analysis: bcbio/bcbio-nextgen. (Blue Collar Bioinformatics, 2019).

19. Friedman, J. H. Greedy Function Approximation: A Gradient Boosting Machine. Ann. Stat. 29, 1189–1232 (2001).

